# Reactive Oxygen Detoxification Contributes to *Mycobacterium abscessus* Antibiotic Survival

**DOI:** 10.1101/2024.10.13.618103

**Authors:** Nicholas A. Bates, Ronald Rodriguez, Rama Drwich, Abigail Ray, Sarah A. Stanley, Bennett H. Penn

**Affiliations:** Department of Internal Medicine, University of California, Davis, California, USA; Graduate Group in Immunology, University of California, Davis, California, USA; Department of Molecular & Cell Biology, University of California, Berkeley, California, USA; Department of Plant & Microbial Biology, University of California, Berkeley, California, USA; Microbiology Graduate Group, University of California, Davis, California, USA; Department of Medical Microbiology and Immunology, University of California, Davis, California, USA

## Abstract

When a population of bacteria is exposed to a bactericidal antibiotic, most cells die rapidly. However, a sub-population of antibiotic-tolerant cells known as “persister cells” can survive for prolonged periods. In addition, antibiotic tolerance can be broadly induced throughout the population by stresses such as nutrient deprivation. However, the pathways required to maintain viability in this setting, and how stress induces antibiotic tolerance are both poorly understood. To identify genetic determinants of antibiotic tolerance in mycobacteria, we carried out transposon mutagenesis insertion sequencing (Tn-Seq) screens in *Mycobacterium abscessus* (*Mabs*) exposed to bactericidal translation-inhibiting antibiotics. This analysis identified genes essential for the survival of both spontaneous persister cells as well as for stress-induced tolerance, allowing the first genetic comparison of these states in mycobacteria. Pathway analysis identified multiple genes involved in the detoxification of reactive oxygen species (ROS), including the catalase-peroxidase *katG*, which contributed to survival in both unstressed and nutrient-starved cells. In addition, we found that endogenous ROS were generated by translation-inhibiting antibiotics, and that hypoxia impaired bacterial killing. *KatG* specifically contributed to survival following exposure to transcription or translation inhibitors, but not other antibiotic classes tested. Thus, the lethality of some antibiotics is amplified by toxic ROS accumulation, and antibiotic-tolerant cells require detoxification systems to remain viable. These findings further demonstrate that antibiotic-induced ROS plays a broad role in mediating antibiotic lethality across diverse organisms.

## INTRODUCTION

A key goal of antibiotic therapy is, in conjunction with the immune system, to eradicate the infecting bacteria. While many common bacterial infections respond rapidly to antibiotics, with 1-2 weeks of therapy sufficient to achieve high cure rates, there are also infections where bacterial clearance is slow and often incomplete^1, 2^. This challenge is exemplified by mycobacterial infections. Fully-susceptible *Mtb* requires multiple antibiotics for four months or longer^3^, and infections by non-tuberculous mycobacteria (NTM), such as *Mycobacterium abscessus* (*Mabs*), are even more difficult to eradicate; *Mabs* often requires treatment for 12-18 months, and even then has a 50% relapse rate^4, 5^.

While the ability of mycobacteria to escape antibiotic-mediated killing is multifactorial, the phenomenon of antibiotic tolerance is likely an important contributor^6–9^. Studies dating from the 1940s noted that when a population of susceptible bacteria were exposed to a bactericidal antibiotic such as penicillin, the majority of the population died within a few hours, but that a small sub-population of cells remained viable for days^10^. Importantly, these antibiotic-tolerant cells referred to as “persister cells” had not acquired a mutation conferring heritable antibiotic resistance, and do not grow in the presence of the antibiotic. Rather, they had entered into a readily reversable phenotypic state where, despite antibiotic-mediated inhibition of critical processes, they were able to survive^11–13^. In addition, a number of physiologic stresses increase antibiotic tolerance in a population, as bacterial cell death is markedly slowed by stresses such as nutrient deprivation or acidic pH^9, 10, 14, 15^. Notably, these same stresses are encountered in the lysosome of an activated immune cell^16^, and studies of pathogens isolated from activated macrophages indeed show a strong immune-mediated increase antibiotic tolerance^8, 17^. Thus, paradoxically, the immune system may actually impede bacterial eradication by antibiotics.

Antibiotic tolerance has been studied extensively in model systems such as *Escherichia coli*, which has provided important insights, but also highlighted uncertainties of current paradigms. Several different regulators of antibiotic tolerance have been identified in *E. coli* including the HipBA toxin-antitoxin system^18^, guanosine pentaphosphate ((p)ppGpp) synthesis by RelA/SpoT enzymes ^19–22^, and Lon protease^23^. In each of these models, the postulated mechanism is to slow metabolism or cell division and render the process targeted by antibiotics non-essential. However, important questions remain. It is unclear how cells remain viable when critical processes, such as transcription or translation, are blocked by antibiotics, as well as how antibiotic tolerance is induced by stress. Even the mechanism of cell death following antibiotic exposure itself has been a matter of debate - originally antibiotics were presumed to kill bacteria as a direct result of inhibition of their target molecule, such as β-lactam antibiotics disrupting cell wall integrity, directly leading to mechanical cell lysis^24^. However, a number of studies, largely from *E. coli*, have demonstrated that in addition to the initial target inhibition, which may be bacteriostatic, bactericidal antibiotics can also cause secondary lethal ROS accumulation, leading to cell death^13,25–30^.

Mycobacterial persister cells are particularly resilient, as *Mycobacterium smegmatis* (*Msmeg*) and *Mycobacterium tuberculosis (Mtb)* persisters can endure many weeks of antibiotic exposure, and stress-induced antibiotic tolerance develops readily^9, 13, 28^. *Mabs* is a species of rapidly-growing mycobacteria, with a doubling time of ∼4h in rich media^31^, and while often environmental, it causes opportunistic infections in patients with structural lung disease such as cystic fibrosis. It is also among the most difficult of all bacterial pathogens to treat, because in addition to forming persister cells, it is also intrinsically resistant to many classes of antibiotics, leaving few treatment options^5^. This leads to the use of antibiotics with greater toxicity to patients and a need to use these agents for prolonged periods to prevent relapse. Thus, identifying the genes that *Mabs* persister cells rely on for survival, as well as the genes involved in the induction and maintenance of stress-induced antibiotic tolerance might highlight pathways that could be targeted therapeutically to eliminate antibiotic-tolerant cells.

Previous genetic screens have studied antibiotic responses in mycobacteria, with some evaluating heritable resistance, and others investigating tolerance. Several studies of resistance have successfully used either transposon mutagenesis with insertion site sequencing (Tn-Seq) or CRISPR-based transcriptional repression with high-throughput sequencing of guide RNAs (CRISPRi) to identify genes promoting growth in sub-inhibitory concentrations of antibiotic. These studies have provided insights, such as highlighting the importance of cell membrane permeability as a mechanism controlling antibiotic penetration into the cytoplasm^32–34^.

Persister formation has proven challenging to study, likely because the low frequency of persisters leads to population bottlenecks that confound genetic analysis. Although screens in *Mtb* have been conducted in macrophages and mice, and genes, such as *glpK* and *cinA* identified, overall the number of mutants isolated in these screens has been low^35, 36^. There has been one effective in vitro Tn-Seq study examining antibiotic tolerance in *Mtb* exposed to starvation and rifampin that isolated over 100 mutants^29^, demonstrating the feasibility of genetic screening in this context. However, whether these phenotypes seen with rifampin in *Mtb* extend to other mycobacteria and other antibiotics remains to be determined.

Here, we study antibiotic tolerance in *Mabs* and describe the results of genome-wide Tn-seq screens seeking to identify the genes required for both spontaneous persister cell survival as well as starvation-induced antibiotic tolerance following exposure to translation-inhibiting antibiotics. We identified several discrete processes contributing to survival and observed a prominent role for ROS detoxifying factors, such as the catalase-peroxidase enzyme KatG, which contributed to both spontaneous persister survival and starvation-induced antibiotic tolerance. Consistent with the protection conferred by KatG, we found that endogenous ROS accumulated following antibiotic exposure, and that the removal of oxygen significantly impaired bacterial killing. Taken together these findings support a model in mycobacteria where the lethality of translation-inhibiting antibiotics is amplified by a secondary accumulation of toxic ROS, and that survival requires active detoxification systems.

## RESULTS

### Starvation-induced antibiotic tolerance in mycobacteria

We first sought to develop conditions suitable for genetic analysis of persister cell survival and stress-induced antibiotic tolerance mycobacteria. Genetic screens examining persister cell physiology face two inherent obstacles. First, these cells are rare in unstressed bacterial populations, and antibiotic-mediated cell death creates population bottlenecks that obscure mutant phenotypes. Second, most mycobacterial populations contain spontaneous drug-resistant mutants that can expand if the population is exposed to a single antibiotic. To overcome these obstacles, we sought to establish large-scale, high-density culture conditions to prevent genetic bottlenecks, and used multiple antibiotics to suppress spontaneous drug-resistant mutants. We began by assessing the feasibility of this approach using wild-type *Msmeg.* We exposed the cells to either the combination of rifampin, isoniazid, and ethambutol (RIF/INH/EMB) used to treat *Mtb*, or to the combination of tigecycline and linezolid (TIG/LZD), two translation-inhibiting antibiotics frequently used to treat *Mabs*^5,37^, and empirically determined the minimum inhibitory concentrations (MICs) and minimum bactericidal concentrations (MBC) for each antibiotic under the high-density culture conditions that would be needed for genetic analysis of persister cells. Both antibiotic combinations reduced the bacterial population >1000-fold within 72h (Figure 1A). We then evaluated both spontaneous persister formation and stress-induced tolerance under these conditions in *Msmeg*. We compared logarithmically growing (mid-log) cultures in 7H9 rich media to cultures starved for 2 days in phosphate buffered saline (PBS) prior to addition of antibiotics. Consistent with expectations, we found a marked increase in antibiotic tolerance in starved cultures, with a 100-fold increase in survival following TIG/LZD exposure and a 10,000-fold increase following RIF/INH/EMB exposure (Figure 1A).

**Figure 1.**
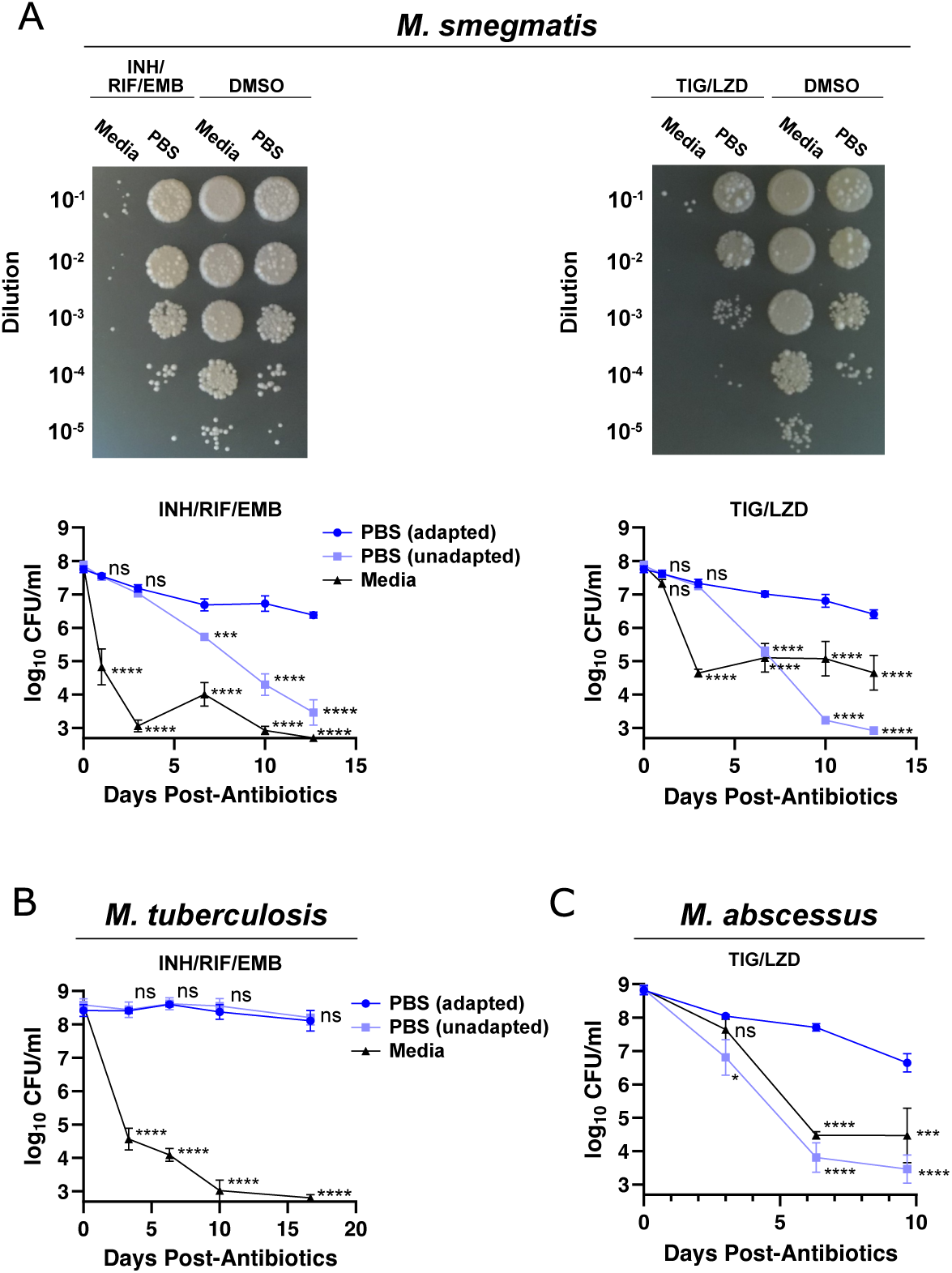
Starvation induces antibiotic tolerance in diverse mycobacteria. (A) *Msmeg,* (B) *Mtb* or (C) *Mabs* were grown in 7H9 rich media or starved in PBS prior to the addition of antibiotics and surviving colony forming units (CFU) enumerated. For the rapidly growing mycobacteria, *Msmeg* and *Mabs,* cells were allowed to adapt for 48h prior to antibiotics, for slow-growing *Mtb,* cells were allowed to adapt for 14-21d prior to antibiotics. Samples without pre-adaptation were washed and placed directly into PBS with antibiotics. Antibiotic concentrations were: *Msmeg -* Isoniazid (INH) 32μg/ml (8 x MIC), rifampin (RIF) 32 μg/ml (8 x MIC), ethambutol (EMB) 4μg/ml (8 x MIC), tigecycline (TIG) 1.25 μg/ml (8 x MIC), linezolid (LZD) 2.5 μg/ml (8 x MIC). *Mtb* – RIF 0.1 μg/ml (4 x MIC), INH at 0.1 μg/ml (4 x MIC), EMB at 8 μg/ml (4 x MIC). *Mabs* – TIG 10 µg/ml (8 x MIC), LZD 100 μg/ml (20 x MIC). Antibiotics with half-lives shorter than the duration of experiment were re-added at the following intervals: TIG, EMB every 3d; RIF, INH every 6d. Error bars represent SEM, statistical significance is calculated at each time point using student’s t test. ****: p<0.0001, ***: p<0.001, **: p<0.01, *: p<0.05, ns: p>0.05. Data are combined from 3 independent experiments.

We next examined two species of pathogenic mycobacteria to assess starvation-induced antibiotic tolerance. Both *Mabs* and *Mtb* have been shown to display this response, and we sought to determine whether this could be observed under the conditions needed to conduct a Tn-Seq screen^38–41^. We again compared cells starved in PBS to logarithmically growing cells in 7H9 and found that, under these conditions, cultures of wild-type *Mabs* (ATCC 19977) and *Mtb* (Erdman) displayed dramatic increases antibiotic tolerance in nutrient-deprived cultures (Figure 1B,C). Notably, for *Mabs* and *Msmeg,* the development of tolerance required an adaptation period of several days under starvation conditions, as survival was dramatically impaired if cells were shifted immediately into nutrient-deficient conditions with antibiotics, suggesting that a regulated process needed to be completed. Surprisingly, *Mtb* tolerance developed rapidly without pre-adaptation suggesting that this organism might have additional response pathways enabling more rapid adaptation.

### Identification of pathways needed for antibiotic tolerance in *Mabs*

We used these conditions to carry out Tn-Seq screens in *Mabs* to identify genes necessary for both the survival of spontaneous persister cells and for starvation-induced antibiotic tolerance. We conducted the screen using a *Mabs Himar1* Tn library comprised of ∼55,000 mutations across ∼91,000 possible TA insertion sites covering all 4,992 non-essential *Mabs* genes in strain ATCC 19977^34^. To study spontaneous persister cells, cultures were maintained in continuous log-phase in 7H9 rich media for 48h prior to antibiotic exposure and then exposed to TIG/LZD for 6 days (Figure 2A) a point at which spontaneous persister cells comprise the majority of the population (Figure 1C). To study starvation-induced tolerance, cultures were starved in PBS for 48h prior to antibiotic exposure in PBS. Following antibiotic treatment, cells were then washed and resuspended in antibiotic-free liquid media to recover and passaged 1:100 three times in continuous log-phase to expand surviving cells. We then isolated genomic DNA, sequenced the Tn insertion sites, and used TRANSIT software^42^ to quantify the abundance of each Tn mutant across different conditions to identify genes with statistically-significant differences in distribution. We identified 277 *Mabs* genes required for surviving TIG/LZD exposure in rich media, 271 genes required for survival during starvation and 362 genes required to survive the combined exposure to antibiotics and starvation (Log_2_ fold-change > 0.5 and Benjamini–Hochberg adjusted p-value (p-adj.) ≤ 0.05; Figure 2B-E). Of the genes required for survival, ∼60% were required in both nutrient-replete and starvation states, although condition-specific determinants were also seen (Figure 2F). As expected, we identified genes with already-established functions in antibiotic responses, including *MAB_2752* and *MAB_2753* which are both homologs of known antibiotic transporters in *Mtb,* as well as tetracycline-responsive transcription factors like *MAB_468*7 and *MAB_0314c* (Supplemental Table 1), indicating an ability of these Tn-Seq conditions to identify physiologically relevant genes known to mitigate antibiotic stress.

**Figure 2.**
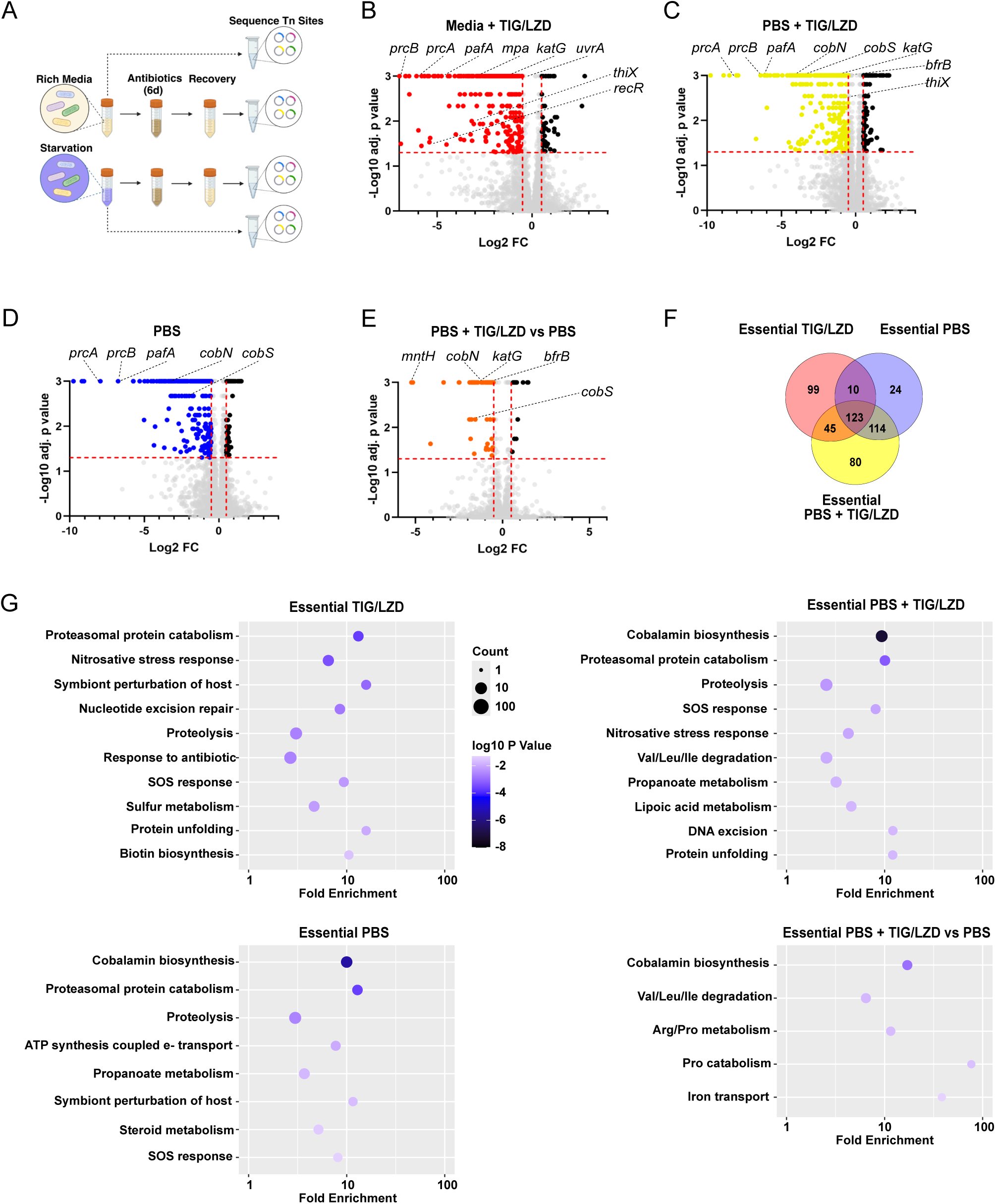
Tn-Seq identifies genes required for antibiotic tolerance in *Mabs*. (A) Experimental design. (B-E) Tn-Seq analysis showing relative abundance of individual genes under the indicted conditions. For (B-D) gene abundance in each condition is measured relative the input with negative values for genes depleted in each experimental condition relative to the input. Log_2_ fold-change is on x-axis with -log_10_ of the p-value on y-axis. All cultures were fully aerated throughout the experiment and cultures without antibiotics received and equal volume of DMSO. In (E) an additional comparison is made for PBS with antibiotics relative to the PBS condition. Genes with significant decreases in abundance are shown in color (p-adj. < 0.05 and log_2_ fold-change > 0.5) using the Benjamini–Hochberg adjustment for multiple hypothesis testing. (F) Number of genes essential in each condition relative to the input population. (G) Pathway enrichment analysis of the essential genes in each condition using the DAVID knowledgebase (p <0.05). Screens were run as 3 independent experiments and the combined results analyzed. Antibiotic conditions were as described above.

To identify other cellular processes necessary for survival we performed pathway enrichment analysis on the set of genes identified by Tn-Seq. We used the DAVID^43^ analysis tool to perform systematic queries of the KEGG, GO, and Uniprot databases and identify over-represented processes and pathways. Interestingly, although cells were exposed to translation-inhibiting antibiotics, and no exogenous oxidative or nitrosative stress was applied, we identified a number of factors needed to combat these stresses in spontaneous persister cells. This included *bfrB* (bactoferritin), *ahpE* (peroxiredoxin) and *katG* (catalase/peroxidase) as well as 5 components of the bacterial proteasome pathway, known to mediate resistance to nitrosative stress in *Mtb*^44^ (Figure 2G, Supplemental Table 2). We also identified multiple members of DNA-damage response pathways including *recF*, *recG*, *uvrA*, *uvrB* and *uvrC*. Examining genes required for starvation-induced tolerance, a number of the same pathways were again seen, and the mutant with the greatest survival defect in this context was *mntH*, a redox-regulated Mn^2+^/Zn^2+^ transporter implicated in peroxide resistance in other organisms^45, 46^.

To independently confirm a role in antibiotic tolerance for a set of genes from diverse pathways that were identified by Tn-Seq we selected a set of genes required for survival, representing several of the functional pathways identified, and used oligonucleotide-mediated recombineering (ORBIT)^47^ to disrupt their open reading frames. The initial genes selected were *pafA* (proteasome pathway), *katG* (catalase-peroxidase), *recR* (DNA repair), *blaR* (β-lactam sensing), and *MAB_1456c* (cobalamin synthesis). To control for non-specific effects of antibiotic selection during the recombineering process, we created a control strain using ORBIT to target a non-coding intergenic region downstream of a redundant tRNA gene (*MAB_t5030c*). We then individually screened each of these mutants to determine if they displayed deficits in survival by exposing cells to TIG/LZD, either in rich 7H9 media or under starvation conditions, as had been done in the Tn-Seq screen. For four out of five mutants we observed defects concordant with the Tn-Seq findings. For *ΔkatG* we detected clear defects in survival as soon as 3 days after antibiotic exposure in either rich media or under starvation conditions, corroborating the results of our Tn-Seq analysis (Figure 3A). We observed similar, albeit smaller, defects in the *ΔpafA, ΔMAB_1456c*, and *ΔblaR,* mutants under the conditions predicted by the screen (Figure 3B-D). We saw no survival defect in the *ΔrecR* mutant (Figure 3E). However, we observed that the *ΔrecR* mutant, as well as the *ΔpafA* mutant, instead displayed a marked defect in resumption of growth after removal of antibiotics (Figure 3 – figure supplement 1).

**Figure 3.**
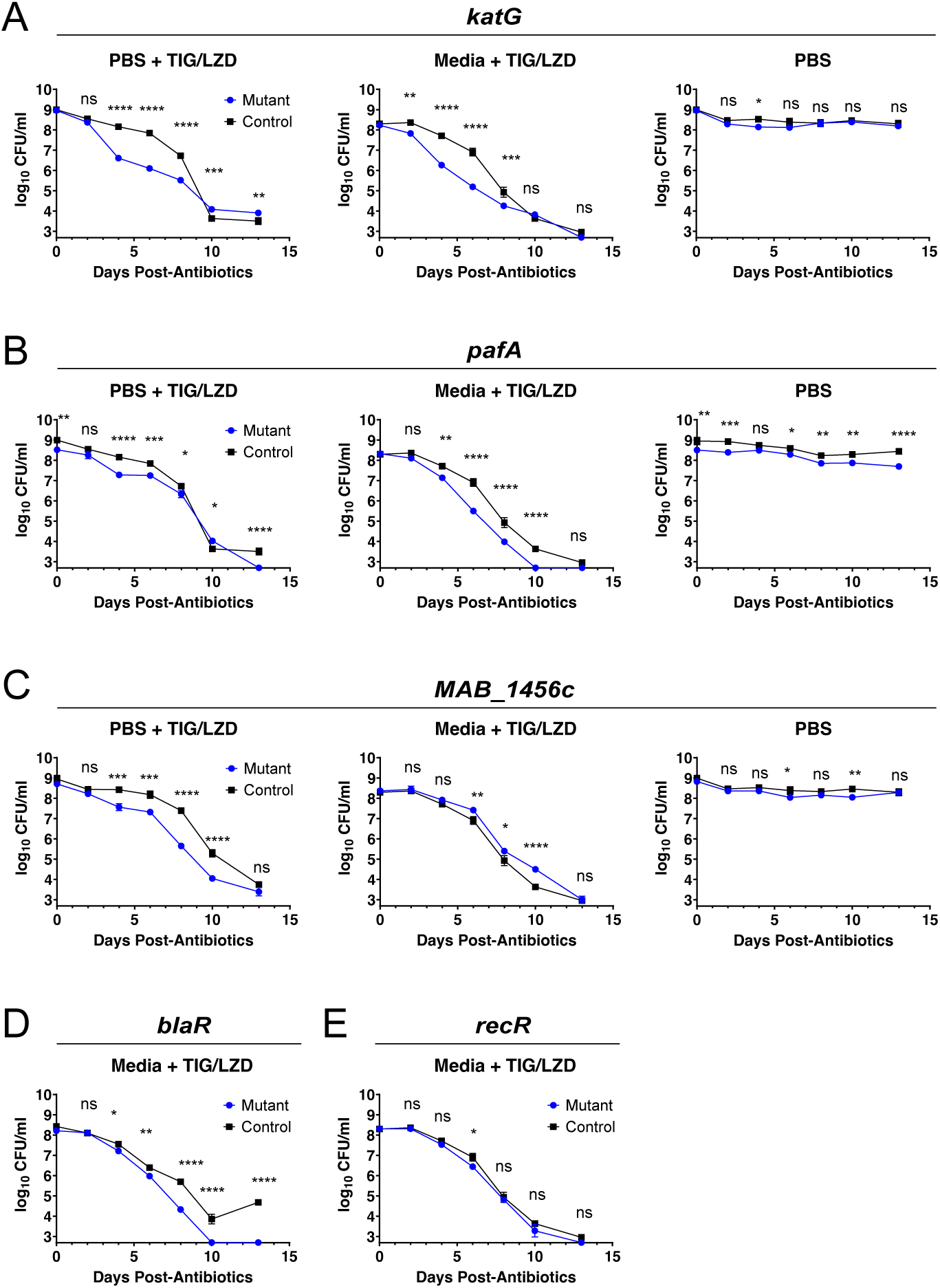
Validation of Tn-Seq results. (A-E) ORBIT homologous recombination was used to delete the indicated genes, or to generate a control strain targeting a distant intergenic region distal to the non-essential tRNA gene *MAB_t5030c*. Each strain was either grown in 7H9 rich media or starved in PBS for 48 prior to the addition of antibiotics as indicated. The conditions tested here correspond to the conditions in the Tn-Seq analysis where a phenotype was observed. Comparisons in panels A-C are made to the same control strain but plotted independently for clarity. Error bars represent SEM, statistical significance is calculated at each time point using student’s t test. ****: p<0.0001, ***: p<0.001, **: p<0.01, *: p<0.05, ns: p>0.05. Antibiotics were added as described above. Data are representative of 4 independent experiments. Figure supplement 1. Additional analysis of mutants

To further confirm the role of *katG* and *pafA,* and exclude off-target effects of recombineering, we performed genetic complementation analysis by restoring a wild-type copy of each gene into the respective *ΔpafA* and *ΔkatG* mutants. In each case, we integrated a single copy of the wild-type gene, under the control of its endogenous promoter, into the genome at the L5 *attB* site (hereafter *pafA+, katG+* strains), and constructed isogenic control strains with an empty vector integrated at the same site (hereafter *pafA-, katG-* strains). We confirmed expression of the re-introduced copy of each gene by RT-qPCR in the *pafA+*, and *katG+* strains, and found expression within roughly 2-fold of endogenous wild-type levels (Figure 4A,D). We then challenged these strains with TIG/LZD as before. In rich media, where the *ΔkatG* mutants have a moderate survival defect, the *katG+* strain had roughly a 50-fold increase in viable cells relative to the *katG-* strain. We then exposed cells to antibiotics under starvation conditions, where the *ΔkatG* mutant phenotype is more severe. Under these conditions the *katG-* cells succumbed rapidly between 3d and 10d after antibiotic exposure, with a 1,000-fold decrease in viable cells relative to control cells, whereas the *katG+* strain showed a near-complete restoration of antibiotic tolerance (Figure 4B). We analogously examined complementation of *ΔpafA* mutants, and although the phenotype of the *ΔpafA* mutant is less severe overall than a *ΔkatG* mutant, we saw a similar restoration of survival in *pafA+* cells relative to *pafA-* cells (Figure 4E). We next evaluated whether the *pafA-*, and *katG-*strains were overall more sensitive to the growth inhibitory effects of TIG/LZD, or whether they had specific defects in survival above the mean bactericidal concentration. We performed MIC determination for TIG and LZD individually for each strain, comparing the *katG+*/*katG-* and *pafA+/pafA-* strains. We found that the MICs for each of these strains were unchanged, demonstrating that these mutants were not more readily inhibited by these antibiotics (Figure 4C,F). Instead, they have more rapid kinetics of cell death at bactericidal concentrations, consistent with a specific defect in survival, and supporting a model whereby an initial growth-arresting inhibition of the direct antibiotic target can be uncoupled mechanistically from a distinct cell-death step, as has been seen with other antibiotic classes^28^.

**Figure 4.**
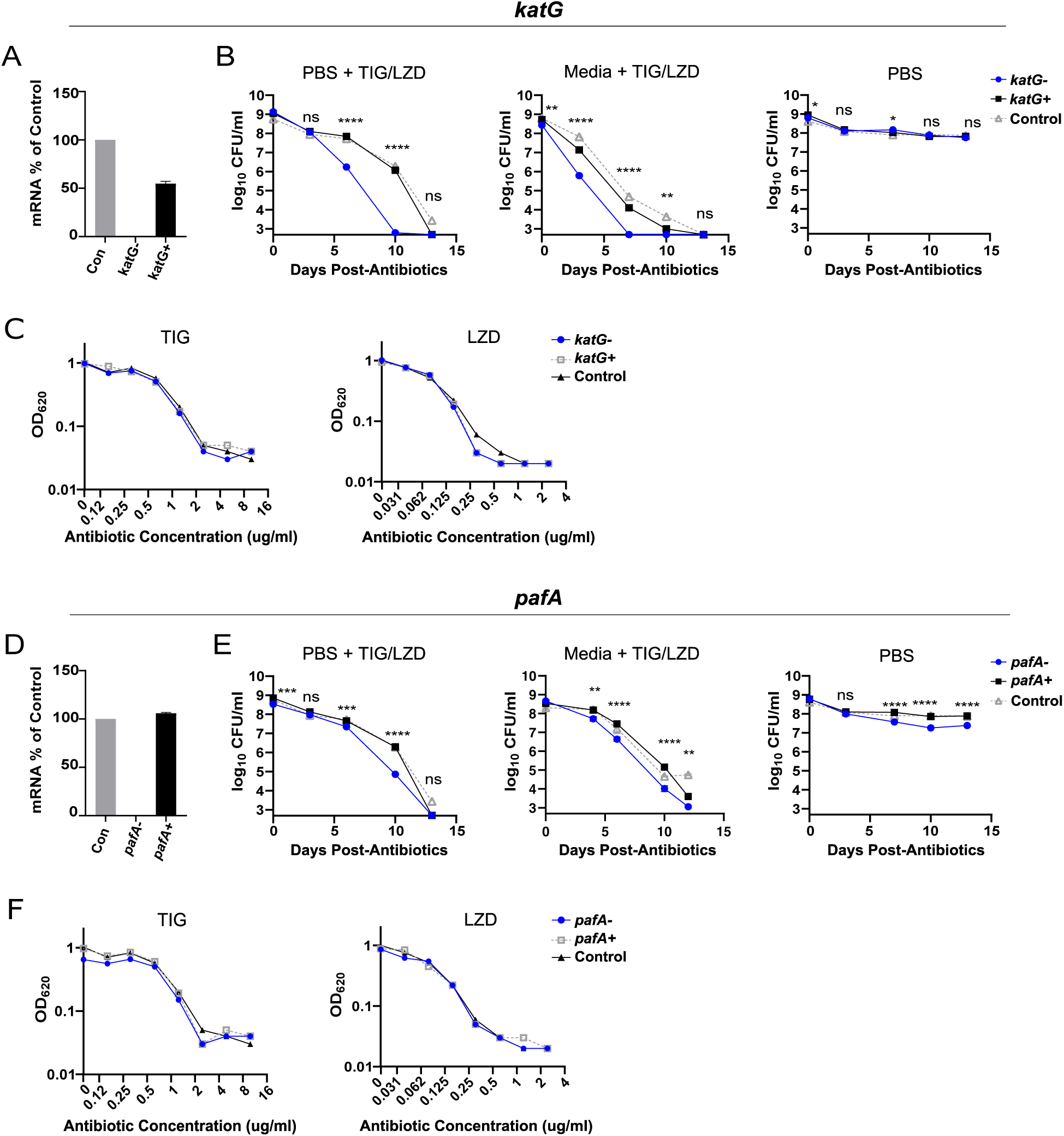
Complementation analysis of *katG* and *pafA* mutants confirms their role in antibiotic tolerance. (A) RT-qPCR analysis of *katG* expression in *katG- (ΔkatG::pmv306), katG+ (ΔkatG::pmv306 katG),* and control strain (ORBIT intergenic*::pmv306*). (B) CFU over time for *katG+/katG-* strains. (C) MICs for *katG+/katG-* strains. (D) Expression of *pafA in pafA-(ΔpafA::pmv306), pafA+* (*pafA::pmv306 pafA*) and control strain. (E) CFU over time for *pafA+/pafA-* strains. (F) MICs for *pafA+/pafA-* strains. Antibiotic concentrations in (A-B, D-E) are as described above. Error bars represent SEM, statistical significance is calculated at each time point using student’s t test between *katG+/katG-* strains in (B) and between *pafA+/pafA-* strains in (E). ****: p<0.0001, ***: p<0.001, **: p<0.01, *: p<0.05, ns: p>0.05. Antibiotics were added as described above.

### Reactive oxygen contributes to antibiotic lethality in *Mabs*

We next investigated the role of KatG and reactive oxygen in antibiotic tolerance more broadly. We began by assessing whether KatG conferred protection from other antibiotics with diverse mechanisms of action, selecting antibiotics that are used clinically for mycobacterial infections. Because *katG-* mutants showed the greatest defects in starvation-induced tolerance, we analyzed survival of *katG+* and *katG-* strains in starvation-adapted cultures exposed to a panel of different antibiotics. Because both TIG and LZD both act by inhibiting translation, we began by exposing cells to either TIG or LZD alone. As expected, the degree of bacterial killing was significantly less with either agent alone than when they are added in combination. Upon exposure to either of these antibiotics the *katG-* cells died more rapidly than *katG+* cells, though the final proportion of persister cells in the population was unchanged in *katG-* cells (Figure 5A). When we exposed cells to rifabutin, an RNA polymerase inhibitor, we saw a similar effect, with a 100-fold loss of viability in *katG-* cells relative to the *katG+* cells (Figure 5B). In contrast, when we exposed cultures to either levofloxacin (topoisomerase inhibitor) or cefoxitin (β-lactam inhibitor of peptidoglycan cross-linking), *katG* had little to no effect on cell viability (Figure 5C-D). Thus, the role of KatG is context-dependent, suggesting that, in *Mabs,* some antibiotics generate oxidative stress that is ameliorated by KatG while others do not.

**Figure 5.**
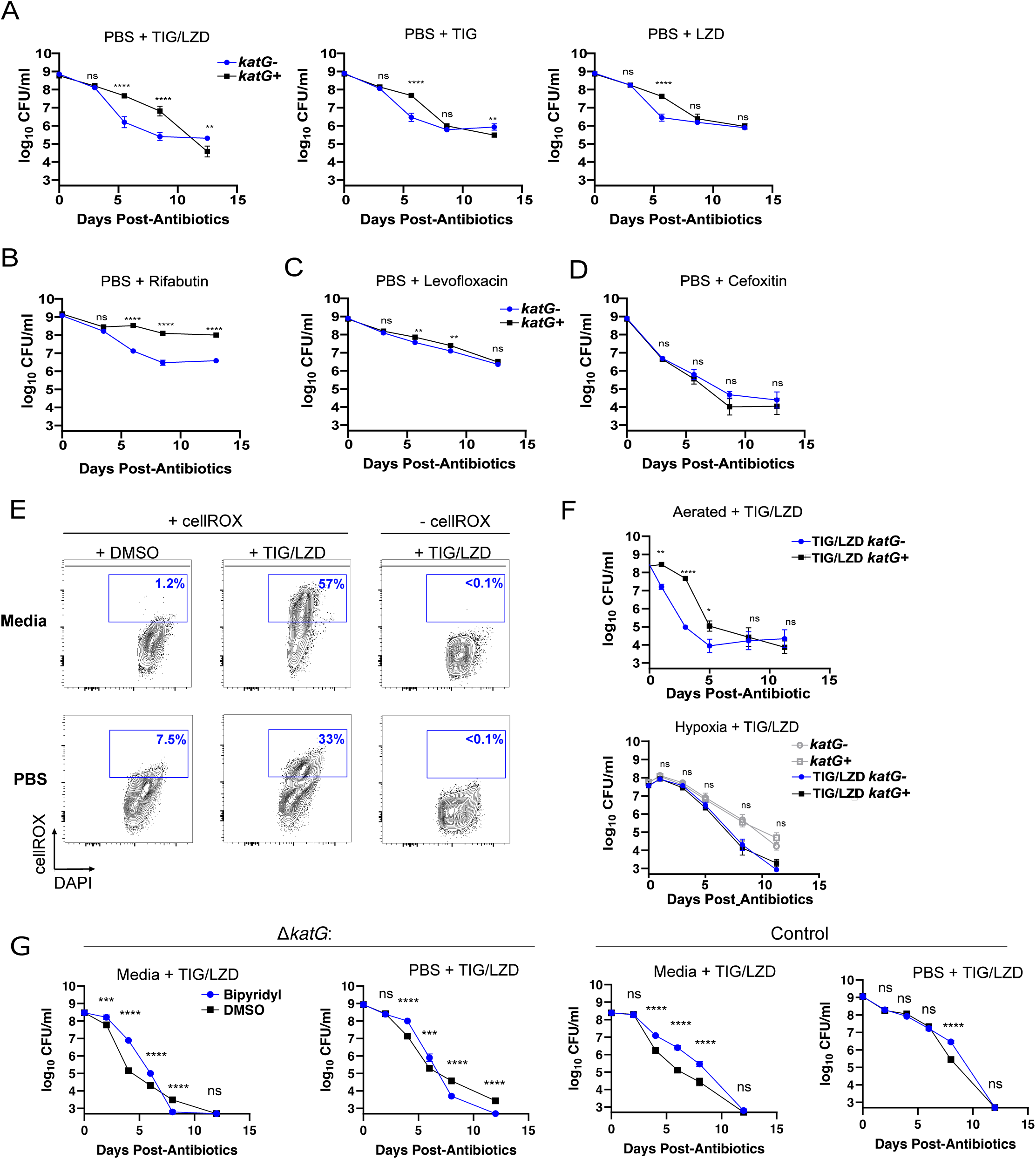
ROS-mediated toxicity following antibiotic exposure. (A-D) Analysis of *katG+/katG-*cells challenged with different antibiotics. Cells were starved in PBS for 48h and then exposed to the indicated antibiotic. (E) Flow cytometry of control cells exposed to TIG/LZD (4h in 7H9 media or 72h in PBS) and then stained with DAPI and the ROS-sensitive dye cellROX green; percentage cellROX-positive cells are indicated. (F) Survival over time for aerated and hypoxic cultures of *Mabs* after exposure to TIG/LZD. (G) Survival over time for bipyridyl treated cells after exposure to TIG/LZD. Error bars represent SEM, statistical significance is calculated at each time point using student’s t test. ****: p<0.0001, ***: p<0.001, **: p<0.01, *: p<0.05, ns: p>0.05. (A-D, F) display combined data from 3 independent experiments. (E, G) are representative data from 3 independent experiments. Figure supplement 1. Effect of ROS scavengers.

The identification of *katG* as essential for cells to survive exposure to TIG/LZD suggests that ROS are present and causing damage. Although TIG and LZD are translation inhibitors that do not directly generate ROS we evaluated whether they might nonetheless be triggering ROS accumulation as a secondary effect. We examined ROS levels in control *Mabs* using the ROS indicator dye cellROX, that is retained in cells when it becomes oxidized^48^. At baseline, during log-phase growth in rich media < 2% of cells had ROS accumulation (Figure 5E). We saw a moderate increase in ROS accumulation in starved cultures, with roughly 7% of the population cellROX+. However, when cells were exposed to antibiotics we saw a dramatic accumulation of ROS, with 57% of cells becoming cellROX+ when exposed to TIG/LZD in rich media and 33% of PBS-starved cells becoming cellROX+ when exposed to TIG/LZD. Taken together, these data indicate that translation inhibition does indeed have important downstream effects on cellular redox balance, with ROS accumulation that could be contributing to the lethal effects of antibiotics.

We next tested whether ROS were contributing to cell death by reducing ROS production, and then assessing the impact on cell viability. A well-established system for studying hypoxia in mycobacteria is the Wayne Model of gradual-onset hypoxia, whereby low density cultures are inoculated in sealed vessels with minimal headspace. As the culture slowly grows, the soluble oxygen is consumed, resulting in the slow onset of hypoxia over several days, a process that can be monitored by the decolorization of Methylene Blue dye in the media^49^. Under aerobic conditions in rich media, we observed the expected rapid killing of *Mabs* over the first 5 days with the combination of TIG/LZD, with more rapid loss of viability in KatG- cells. However, under hypoxic conditions, where ROS production is suppressed, we saw much slower bacterial killing.

Importantly, under hypoxic conditions *katG-* cells no longer had a survival defect relative to *katG+* cells, supporting the hypothesis that translation-inhibiting antibiotics also cause secondary accumulation of lethal ROS in antibiotic-treated cells that need to be detoxified by KatG (Figure 5F).

We also evaluated whether other methods of alleviating ROS damage might enhance survival. The iron chelator 2,2’-dipyridyl has been shown in other contexts to reduce ROS-mediated damage by suppressing the reaction of H_2_0_2_ with Fe^2+^ that generates highly oxidizing hydroxyl radicals (Fenton reaction), and which has been shown to mitigate oxidative damage in other bacteria following antibiotic exposure^25, 28, 50^. As seen in other bacteria, we find that in *Mabs*, 2,2’-dipyridy does indeed improve bacterial survival following exposure to bactericidal translation inhibitors, further supporting a role in ROS in cell death following translation inhibition (Figure 5G).

We also assessed whether free radical scavengers like thiourea and 2,2,6,6-tetramethylpiperidine-1-oxyl (TEMPO) ameliorated antibiotic toxicity, although similar thiol antioxidants had previously been shown in *Mtb* to increase respiration and ROS generation and to paradoxically decrease bacterial survival upon INH exposure^30^. When we treated *Mabs* simultaneously with TIG/LZD in combination with either thiourea or TEMPO we did not observe a restoration of antibiotic tolerance, and, similar to observations in *Mtb,* actually observed increased bacterial cell death (Figure 5 – figure supplement 1).

### Antibiotic-induced ROS accumulation is conserved but reliance on KatG is variable among *Mabs* strains

To test whether ROS accumulation was an effect occurring more broadly across different *Mabs* strains we obtained two clinical strains, exposed them to TIG/LZD in 7H9 media, and measured ROS accumulation with cellROX as above. We found that similar to ATCC 19977, both clinical

*Mabs* strains had elevated ROS levels following translation inhibition (Figure 6A), suggesting that this is a conserved process in *Mabs*. Next, we tested the role of KatG in these *Mabs* clinical strains. We used ORBIT to disrupt the *katG* locus and evaluated the ability of these *ΔkatG* clinical strains to survive exposure to TIG/LZD under both stressed and un-stressed conditions. Unlike ROS accumulation, where the responses across strains were consistent, we saw a variable dependency on *katG*. Clinical strain-1 behaved differently overall, with no appreciable starvation-induced antibiotic tolerance, and no contribution of *katG* to survival. In contrast, for clinical strain-2, *katG* contributed significantly to starvation-induced antibiotic tolerance, behaving similarly to the ATCC 19977 reference strain. However, unlike the reference strain, *katG* was not required for survival of clinical strain-2 when exposed to antibiotics in 7H9 media (Figure 6B). Thus, although antibiotic-induced ROS accumulation was observed across all three *Mabs* strains, the *ΔkatG* phenotype displays incomplete penetrance, suggesting that in some *Mabs* strains alternative pathways exist that are able to compensate for the loss of *katG*.

**Figure 6.**
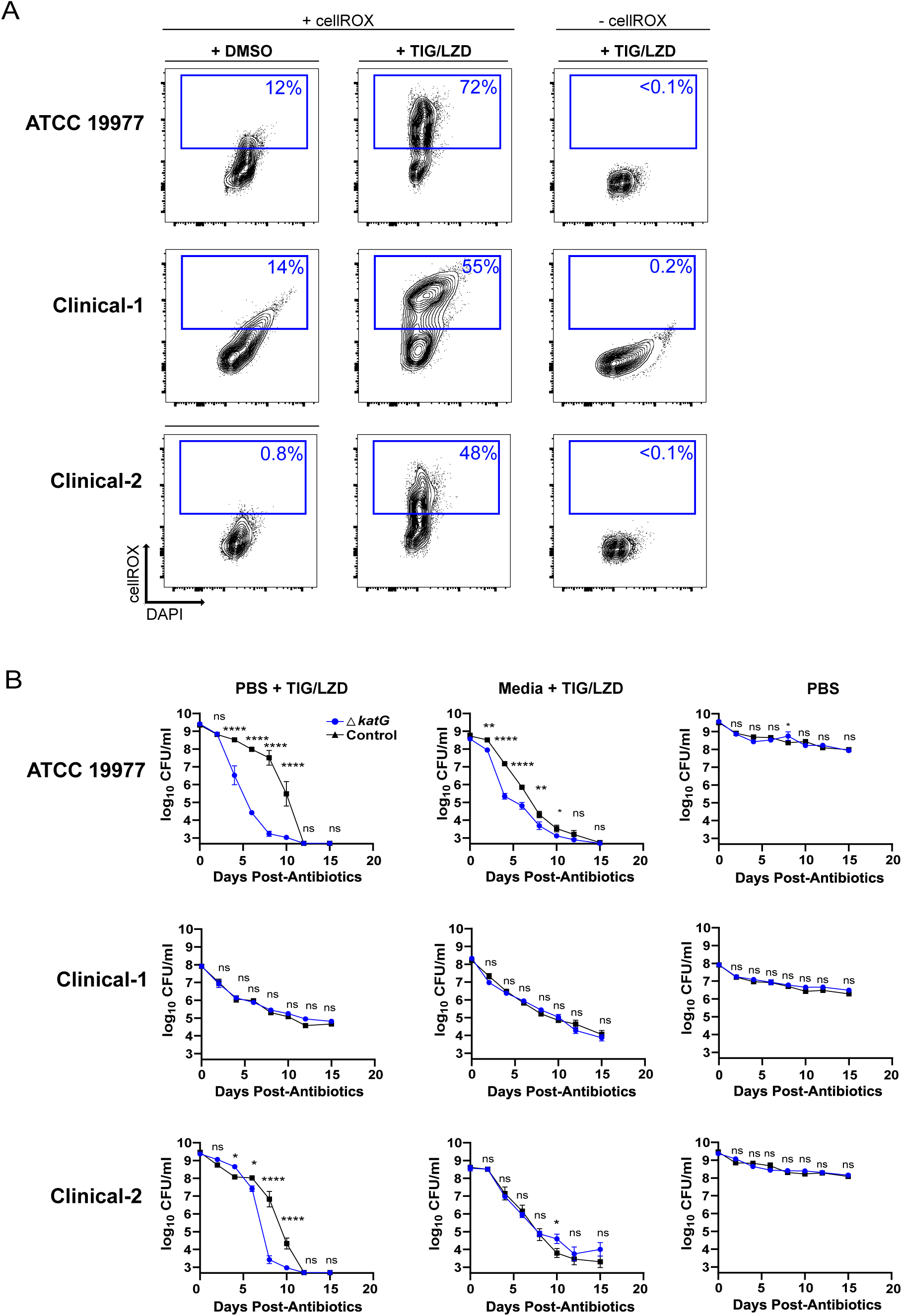
Conserved antibiotic-induced ROS production but variable protection by KatG among different *Mabs* strains. (A) The indicated strains of *Mabs* were cultured in 7H9 media, exposed to TIG/LZD for 4h and then analyzed by cellROX staining. (B) CFU over time following TIG/LZD exposure. Error bars represent SEM, statistical significance is calculated at each time point using student’s t test. ****: p<0.0001, ***: p<0.001, **: p<0.01, *: p<0.05, ns: p>0.05. Combined data from 4 independent experiments are shown for persister survival experiments. Representative data from 2 independent experiments are shown for flow cytometry experiments. Antibiotics were added as described above.

## DISCUSSION

The results of these studies point to an important effect of ROS in amplifying the lethality of transcription and translation-inhibiting antibiotics in *Mabs*. Through genetic analysis we identified a number of ROS detoxification factors, including KatG, as necessary for survival in this context. This suggested that antibiotics induced an oxidative state in cells, and direct measurement of ROS following antibiotic exposure indicated that this was indeed the case. Further supporting the toxic effects of ROS in this context, we found that removal of oxygen both slowed bacterial killing and rendered KatG dispensable. Taken together, these results suggest that in *Mabs* antibiotic lethality is accelerated by ROS accumulation, and that survival requires active detoxification systems.

### Pathways necessary for antibiotic tolerance in *Mabs*

The phenomenon of antibiotic tolerance has been recognized for decades and has been observed in a broad array of bacterial species^10, 51–57^ but without identification of a singular underlying mechanism conserved among species - suggesting that different pathways may play roles in different physiologic contexts. For example, the pathways identified in one bacterial species may not contribute to tolerance in another. In *E. coli, relA* plays an important role in stress-induced tolerance, as it does in *Pseudomonas aeruginosa*^52^*, Staphylococcus aureus*^53^, and *Mtb*^54^. However, its role is not universal. Deletion of *relA* had no effect on antibiotic tolerance in *Msmeg*^55^, and in our Tn-Seq analysis, *Mabs relA* Tn mutants had no survival defect. In the case of *Mabs* this may be due to genetic redundancy, as a prior study of the *Mabs relA* mutant demonstrated that this strain still synthesizes (p)ppGpp^58^. In addition, even within a single species there can be differences in the critical survival mechanisms depending on the context. In *E. coli,* RelA contributes strongly to persister viability following exposure to β-lactams, but not aminoglycosides^56^, and in our study we find KatG to be essential for tolerance to transcription and translation inhibitors but not to a β-lactam (cefoxitin) or a quinolone (levofloxacin).

### Mechanisms of antibiotic lethality

Our findings strongly support the idea that antibiotic-induced ROS can be a significant contributor to bactericidal activity in *Mabs* and contribute to the growing evidence that this phenomenon is conserved across diverse types of bacteria. In mycobacteria, other groups have observed that hypoxia reduced antibiotic-mediated killing in *Mabs, Mtb* and *Msmeg*, and in *Mtb*, exposure to rifampin or moxifloxacin also generates ROS, with *katG* contributing to survival in rifampin-treated cells^28, 29^. Notably, we found evidence for ROS mediated bactericidal activity with the translation inhibiting antibiotics tigecycline and linezolid, as well as the transcription inhibiting antibiotic rifabutin. This contrasts with findings in *E. coli* where the rifamycins and tetracyclines are not bactericidal and do not induce ROS^25, 26^, suggesting that mycobacteria-specific responses may exist that result in lethal ROS production.

Exactly how transcription or translation blockade leads to increased ROS levels is not known. In principle, any of several derangements could lead to ROS accumulation. One of the major sources of cellular ROS is oxidative phosphorylation, as hydrogen peroxide and superoxide are natural byproducts. Thus, increased ROS generation by oxidative phosphorylation is an attractive hypothesis. Alternatively, particularly under starvation conditions, it is possible that antioxidants and ROS scavengers may become depleted, creating a more oxidizing environment. Our Tn-Seq analysis provides additional insight on this. We noted a small class of Tn mutants that were paradoxically protected from antibiotic lethality (Figure 2B). Prominent among this class of mutants were several independent components of the NADH dehydrogenase complex. Also known as Complex I of the electron transport chain, it is one of the key entry points for electrons into the oxidative phosphorylation pathway. The observation that mutants lacking this complex are protected suggests that decreasing flux through oxidative phosphorylation, with a concomitant decrease in ROS generation, may enhance survival during antibiotic exposure. A mechanistic understanding of how blockade of either transcription or translation leads to deranged oxygen utilization is an unresolved question that will require further study.

ROS accumulation is not universal following exposure to bactericidal antibiotics. While antibiotic-induced ROS is well-documented, under some conditions, such as higher concentrations of antibiotic, multiple studies have also observed antibiotic lethality without ROS accumulation^59–62^. Similarly, prior studies have found that under certain conditions, *E. coli* mutants lacking catalase have defects in antibiotic tolerance^63–65^, whereas under other conditions they do not^59^ . In *Mabs* the role of *katG* was also not uniform, as it had no impact on survival following exposure to levofloxacin or cefoxitin. Additional studies will be needed to determine whether levofloxacin and cefoxitin kill without generating ROS, or whether ROS is generated but effectively detoxified by other systems in the absence of *katG.* This latter possibility is suggested by our findings in *Mabs* clinical strains. While we saw ROS accumulation in all strains after TIG/LZD exposure, the role of *katG* was variable between the strains. This suggests that compensatory pathways likely exist and that in *Mabs* they can overcome the loss of *katG*.

### Therapeutic implications

*Mabs* infections are particularly challenging to treat, and frequently have poor outcomes^4^. Our results highlight several bacterial processes, such as the bacterial proteasome and ROS detoxification, which might be targeted therapeutically to reduce the survival of antibiotic tolerant bacteria in patients with *Mabs* infection. Agents targeting these processes might not have any intrinsic antimicrobial activity alone, but might act to disrupt the unique physiology required to survive antibiotics. This would represent a new therapeutic class of “persistence inhibitors” that might act synergistically with traditional antibiotics to eliminate the subpopulation of cells that would otherwise remain viable, despite prolonged antibiotic treatment, in patients with *Mabs* and other chronic infections^28, 30^.

### Limitations

Tn-Seq has inherent drawbacks, including an inability to identify mutants in essential genes, or in cases of genetic redundancy. Thus, there are likely genes needed for antibiotic tolerance in *Mabs* that were not identified in this study. In addition, we studied the response to a single class of antibiotic, focusing on the translation inhibitors often used to treat *Mabs* infections, and we studied only spontaneous persister cells and starvation-induced antibiotic tolerance. It is likely that examining other antibiotics, with different mechanisms of action, or different stresses that induce tolerance would identify additional genes contributing to survival and would allow identification of core pathways that might be shared in differing physiologic contexts of antibiotic and stress.

## MATERIALS AND METHODS

### Key Reagent Table

| **Reagent** | **Source/Reference** | **Identifier** |
| --- | --- | --- |
| ***Mabs* ATCC 19977** | ATCC | 19977 |
| ***Msmeg* MC2 155** | ATCC | 700084 |
| ***Mtb* Erdman** | ATCC | 35801 |
| ***Mabs* clinical strains** | Sacramento Department of<br>Public Health | -- |
| **pkm444** | Addgene | 108319 |
| **pkm496** | Addgene | 109301 |
| **pmv306** | Ref.66 | -- |
| **Middlebrook 7H9** | BD | 271310 |
| **Middlebrook 7H10** | BD | 262710 |
| **Glycerol** | Invitrogen | 15514011 |
| **Tween-80** | Fisher | BP338-500 |
| **Middlebrook OADC** | BD | 212351 |
| **DPBS (-Ca/Mg)** | Gibco | 14190144 |
| **Tyloxapol** | Sigma Aldrich | T8761 |
| **Methylene blue** | Sigma Aldrich | 28514 |
| **Tigecycline** | Chem Impex | 29737 |
| **Linezolid** | Chem Impex | 29723 |
| **Levofloxacin** | Sigma Aldrich | 28266 |
| **Cefoxitin** | Chem Impex | 1490 |
| **Rifabutin** | Cayman Chemical | 16468 |
| **Rifampin** | Sigma Aldrich | R7382 |
| **Isoniazid** | Supelco | I3377 |
| **Ethambutol** | Thermo Scientific | J6069506 |
| **DMSO** | Sigma Aldrich | D2650 |
| **Trizol** | Invitrogen | 15596018 |
| **0.1mm zirconia beads** | Biospec | 11079101z |
| **PureLink RNA Mini Kit** | Invitrogen | 12183025 |
| **Kanamycin** | VWR | 75856-686 |
| **Zeocin** | Invivogen | ANTZN1P |
| **Glycine** | Fisher Bioreagents | BP381 |
| **Sucrose** | Sigma Aldrich | S1888 |
| **Anhydrotetracycline** | Cayman Chemical | 10009542 |
| **cellROX green** | Invitrogen | C10444 |
| **Paraformaldehyde** | Sigma Aldrich | P6148 |
| **DAPI** | Sigma Aldrich | D9542 |
| **DNAseI** | New England Biolabs | M0303 |
| **Maxima H minus reverse transcriptase** | Thermo Scientific | EP0752 |
| **Taq Polymerase** | New England Biolabs | M0273 |
| **EvaGreen** | Biotium | 31019 |
| **2,2'-Bipyridyl - 2,2'-Bipyridine** | Sigma Aldrich | D216305 |
| **Thiourea** | Sigma Aldrich | T7875 |
| **4-Hydroxy-TEMPO** | Sigma Aldrich | 176141 |

#### Bacterial strains and culture conditions

*Mabs* ATCC 19977, clinical *Mabs* strains and *Msmeg* (MC2 155) were grown in BD Middlebrook 7H9 media (liquid) or 7H10 media (solid) supplemented with 0.5% glycerol (Sigma) and 0.2% Tween-80 (Fisher) but without any OADC supplementation except for transformations. Sacramento clinical isolates were obtained from the Sacramento County Department of Public Health Mycobacteriology Laboratory. Confirmation of clinical isolates as *Mabs* was performed by amplifying the 16s rRNA locus and Sanger sequencing (Supplemental Table 4). *Mtb* (Erdman) was grown in 7H9 (liquid) or 7H10 (solid) supplemented with 0.5% glycerol, 0.1% Tween-80, and 10% OADC (BD). All cultures were grown at 37°C with gentle shaking. Except for specific hypoxia conditions, all liquid cultures were grown with 90% container headspace or using a gas permeable cap to ensure culture oxygenation. PBS starvation was achieved by washing OD 0.5-1.0 *Mabs* 1X in DPBS (-Ca/Mg, Gibco) and resuspending in DPBS at OD=1, supplemented with 0.1% tyloxapol (Sigma).

#### *Mabs* antibiotic experiments

For PBS starvation experiments, stocks of *Mabs* were grown for 48 hours in 7H9 passaging continuously in log phase, then either PBS starved or passaged in log phase for an additional 48 h. Log phase or PBS starved *Mabs* were then resuspended in antibiotic containing media at OD1.0. For experiments with hypoxia, *Mabs* in mid-log aerobic growth was adjusted to OD 0.001 in media with 1.5 μg/ml methylene blue and added to a rubber septum sealed glass vial with 50% headspace. Methylene blue discoloration was observed at 3d and antibiotics were added at 5d. For thiourea and TEMPO experiments, these were added to the cultures at the time of antibiotic administration. For bipyridyl experiments, 62.5 μM bipyridyl was added to the cultures 2h prior to antibiotic administration. We empirically determined the half-life of each antibiotic in 7H9 media at 37-deg and for those with half-lives shorter than the experiment, supplemented cultures to with additional antibiotic to maintain the concentration of active antibiotic. Antibiotics were used at the following concentrations: tigecycline (Chem-Impex) at 10 μg/ml (8-fold above MIC, re-administered every 3 days), linezolid (Chem-Impex) at 100 μg/ml (20-fold above MIC), levofloxacin (Sigma) at 40μg/ml (8-fold above MIC), cefoxitin (Chem Impex) at 80 μg/ml (8-fold above MIC, re-administered every 3 days), and rifabutin (Cayman) at 40μg/ml (4-fold above MIC). After antibiotic administration, colony forming units over time were measured by plating on 7H10 solid media. For experiments where growth recovery time in liquid media was quantified, 100μl of sample was removed at d6 after antibiotic administration and washed 2X in antibiotic-free media. The samples were resuspended in 5ml of antibiotic free media, and OD620 measurements were taken with a FilterMax F3 plate reader (Molecular Devices) until maximum cell density (OD of ∼5.0) was reached.

#### *Msmeg* antibiotic experiments

Individual colonies were picked and grown for 48 hours in log phase before being PBS starved or passaged in log phase for 48h. Log phase or PBS starved *Msmeg* were then resuspended in antibiotic containing media at OD1.0. Antibiotics were used at the following concentrations: tigecycline (Chem-Impex) at 1.25 μg/ml (8-fold above MIC, re-administered every 3 days), linezolid (Chem-Impex) at 2.5 μg/ml (8 fold above MIC), rifampin (Sigma) at 32 μg/ml (8-fold above MIC, re-administered every 6 days), isoniazid (Sigma) at 32 μg/ml (8-fold above MIC, re-administered every 6 days), and ethambutol (Thermo) at 4 μg/ml (8-fold above MIC, re-administered every 3 days). After antibiotic administration, colony forming units over time were measured.

#### *Mtb* antibiotic experiments

Freezer stocks of *Mtb* were thawed and grown for 5-7 days in log phase before being starved for 14d or longer. Non-starved control *Mtb* were thawed such that they were also grown for 5-7 days in log phase before experimental use. Log phase or PBS starved *Mtb* was then resuspended in antibiotic containing media and adjusted to OD 1.0. Antibiotics were used at the following concentrations: rifampin (Sigma) at 0.1 μg/ml (4-fold above MIC, re-administered every 6 days), isoniazid (Sigma) at 0.1μg/ml (4-fold above MIC, re-administered every 6 days), and ethambutol (Thermo) at 8 μg/ml (4-fold above MIC, re-administered every 6 days). After antibiotic administration, colony forming units over time were measured.

#### Transposon insertion sequencing

The construction of this *Himar1* transposon Tn library has been described previously^34^. Screening was performed by growing a freezer stock of the library for 2.5 days in log phase before 48 h PBS starvation or further continuous log-phase growth. Samples were then resuspended in media containing either tigecycline/linezolid or an equal volume of DMSO solvent and incubated for 6 days, with a re-administration of tigecycline or matching DMSO on day 3. Cultures were aerated by culturing in a vented cap bottle with gentle agitation at 40 revolutions per minute throughout the experiment. The samples were then washed 2X in antibiotic free liquid media, resuspended in antibiotic free liquid media (10X the original culture volume), and grown until OD=0.5-1.0. Subsequently, the samples underwent three more rounds of 100-fold passaging in liquid media to amplify surviving bacteria before the samples were collected in Trizol (Invitrogen). A sample taken at the time of the commencement of PBS starvation was collected in Trizol and used as the input control. Three independent trials of this experiment were submitted to the UC Davis DNA Technologies Core, where Tn insertion site flanking sequences were amplified as described previously^34^ and sequenced on an Element Biosciences AVITI. Sequence reads were mapped to the ATCC 19977 genome and analyzed using TRANSIT software with the following parameters: 0% of N/C termini ignored, 10,000 samples, TTR normalization, LOESS correction, include sites with all zeros, site restricted resampling. Genes with significant changes were defined as those with adjusted p-value (p-adj.) <0.05 and log_2_ fold change >0.5. P-adj. was calculated using the Benjamini-Hochberg correction.

#### Pathway enrichment analysis

To improve gene annotation, *Mabs* orthologs to *Mtb* genes were identified. *Mabs* genes were first converted into protein sequences using Mycobrowser, and protein sequences were then used to perform reciprocal BLASTp searches. *Mabs* genes and *Mtb* genes that mapped to each other using independent one-way BLASTp searches with a maximum e-value cutoff of 0.1 were considered orthologs. For pathway analysis, gene lists (*Mtb* orthologs) were then imported into the DAVID knowledgebase^43^ and pathway enrichment analysis performed for Gene Ontology biological process, Uniprot keyword and KEGG databases with statistical analysis Fisher’s exact test and nominal p-value reported.

#### Gene deletion and complementation

Knockout strains were generated using ORBIT^47^. Briefly, *Mabs* was transformed with the kanamycin-resistant ORBIT recombineering plasmid pkm444. 20ml *Mabs* at OD=0.5-1.0 was washed 2X in 10% glycerol and resuspended in 200ul 10% glycerol. 500ng plasmid was added and electroporated at 2.5kV in 0.2cm cuvettes. The bacteria were allowed to recover overnight before plating on 150μg/ml kanamycin plates. Clones were selected and regrown in liquid media supplemented with 150μg/ml kanamycin and 10% OADC (BD) to OD0.5-1.0. For recombineering, the pkm444-*Mabs* was grown to mid-log, then diluted to OD0.1 and 200mM glycine (Fisher) was added to the media. 16 h later, 500mM sucrose (Sigma) and 500 ng/ml anhydrotetracycline (Cayman) were added and incubated for an additional 4 h. Subsequently, the *Mabs* was washed 2X in ice cold 10% glycerol + 500mM sucrose. 200ul of 10X concentrated *Mabs* was then electroporated with 600ng of the zeocin-R ORBIT payload pkm496 plasmid and 2μg of targeting oligonucleotide (Table S3) at 2.5kV in 0.2cm cuvettes. The *Mabs* was then allowed to recover overnight in liquid media with 10% OADC and 500ng/ml anhydrotetracycline before being plated on 150 μg/ml zeocin plates. Mutants were then selected and screened for gene deletion by PCR amplification and Sanger sequencing. For genetic complementation, the endogenous loci including promoter and terminator sequences were amplified by PCR and cloned into the EcoRV site of pmv306 with kanamycin resistance^66^. In the case of *katG,* the upstream gene *furA* was also included in the complementation construct to achieve optimal *katG* expression.

#### MIC determination

Two-fold serial dilutions of antibiotics were prepared in a 96 well plate in 100ul volume. 100ul of 2X bacteria were added (for *Mabs*: used a final OD of 0.001, *Msmeg*: OD0.001, *Mtb*: OD0.01), making a final volume of 200μl. The plates were incubated until there was visible growth in the no antibiotic control well. At this time, the bacteria were transferred to a new plate with 20μl of 40% paraformaldehyde and OD620 measurements were taken with a FilterMax F3 plate reader (Molecular Devices). MIC values for wild-type ATCC 19977 under these conditions are listed in Supplemental Table 5.

#### Flow cytometry

A culture of OD=1.0 *Mabs* was stained with cellROX green (Invitrogen) at a final concentration of 5μM for 1hr at 37C. The cells were then washed in PBS and resuspended in PBS with 4% paraformaldehyde and 5 μg/ml DAPI (Sigma). The samples were run on an LSRII flow cytometer (BD). Fluorophores were excited with the 405nm (DAPI) and 488nm (cellROX) lasers. Detection was performed using the 450/50 (505LP) filter for DAPI and a 525/50 (555LP) filter for cellROX. Data were analyzed with FlowJo software (BD).

#### DNA/RNA Purification

Samples were resuspended in 5 volumes of Trizol, and bead beat with 0.1mm zirconia beads (Biospec) 6x2min at 4°C in a Mini-Beadbeater-16 (Biospec). Chloroform was added and RNA in the aqueous phase removed. For DNA isolation, a second RNA extraction was performed with 0.8M guanidine thiocyanate and 0.5M guanidine hydrochloride, 60mM Acetate pH 5.2, 1mM EDTA. DNA was then isolated with back-extraction buffer (4M Guanidine Thiocyanate, 50mM Sodium Citrate, 1M Tris base (without pH adjustment ∼pH 11) and DNA purified using a PureLink RNA Mini Kit (Invitrogen).

#### RT-qPCR

RNA was purified using PureLink RNA Mini Kit per manufacturer’s instructions. The samples were DNAseI (NEB) treated for 15min/37°C before stopping the reaction by adding 3.5mM EDTA and heating for 10min/75°C. cDNA was synthesized from 500ng total RNA using random hexamers and Maxima H minus reverse transcriptase (Thermo). No reverse-transcription controls were also included and used to confirm the lack of genomic DNA-driven amplification. qPCR reactions used Taq polymerase (NEB) and EvaGreen (Biotium) and were run on Biorad CFX Opus 96 Real-Time PCR System. Melt curves were included for each sample to confirm uniform amplicon identity between samples. Gene-specific amplification was quantified by comparison to a standard curve generated from 3-fold serial dilutions of a control sample, then normalized to 16S rRNA within each sample.

## ACKNOWLEDGEMENTS

We would like to thank Nick Campbell-Kruger for his technical insights on identifying *Mabs/Mtb* orthologs, Jonathan Van Dyke for his assistance with flow cytometry analysis, Emily Kumimoto and Siranoosh Ashtari for their assistance with Tn-Seq library preparation and sequencing, and Jessie Li and Bradley Jenner for their assistance with Tn-seq bioinformatics. We would also like to thank Caroline Dominic at the Sacramento County Department of Public Health for providing clinical isolates of *Mabs* for analysis.

## FUNDING

Pew Biomedical Scholars Fellowship BHP NIH R01 1R01AI144149 BHP NIH R01 1R01AI143722 SAS NIH T32 HL007013 AR

NIH Shared Instrumentation Grant 1S10OD010786-01 DNA Technologies and Expression Analysis Core, UC Davis Genome Center

NCI Cancer Center Support Grant P30CA093373 Flow Cytometry Shared Resource, UC Davis

## CONFLICTS OF INTEREST

BHP and SAS serve on the scientific advisory board of X-Biotics Therapeutics.

---

**Figure 3 – figure supplement 1.**
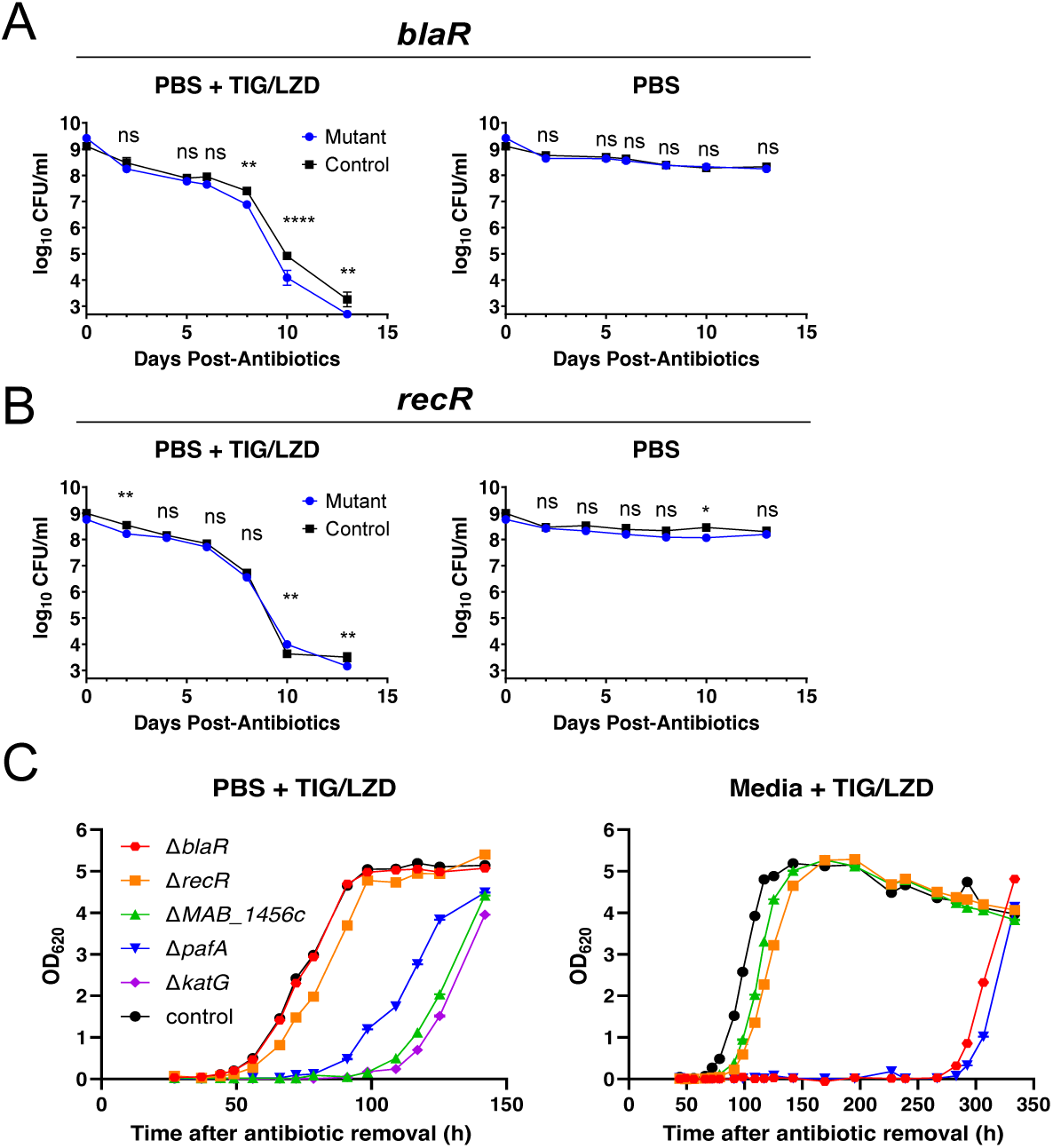
Additional analysis of mutants. (A) *blaR* or (B) *recR* mutants were examined under conditions where Tn-Seq did not predict a phenotype. Cells were starved in PBS for 48h prior to treatment with TIG/LZD or DMSO as described in Figure 3. (C) Growth recovery of mutants after antibiotic exposure. After 6 d of TIG/LZD exposure cells were washed twice in antibiotic free media and inoculated into 7H9 media. Growth was monitored by OD 600 of the culture. Error bars represent SEM, statistical significance is calculated at each time point using student’s t test. ****: p<0.0001, ***: p<0.001, **: p<0.01, *: p<0.05, ns: p>0.05. (A-B) are representative data from 4 independent experiments (C) is representative data from 2 independent experiments.

**Figure 5 – figure supplement 1.**
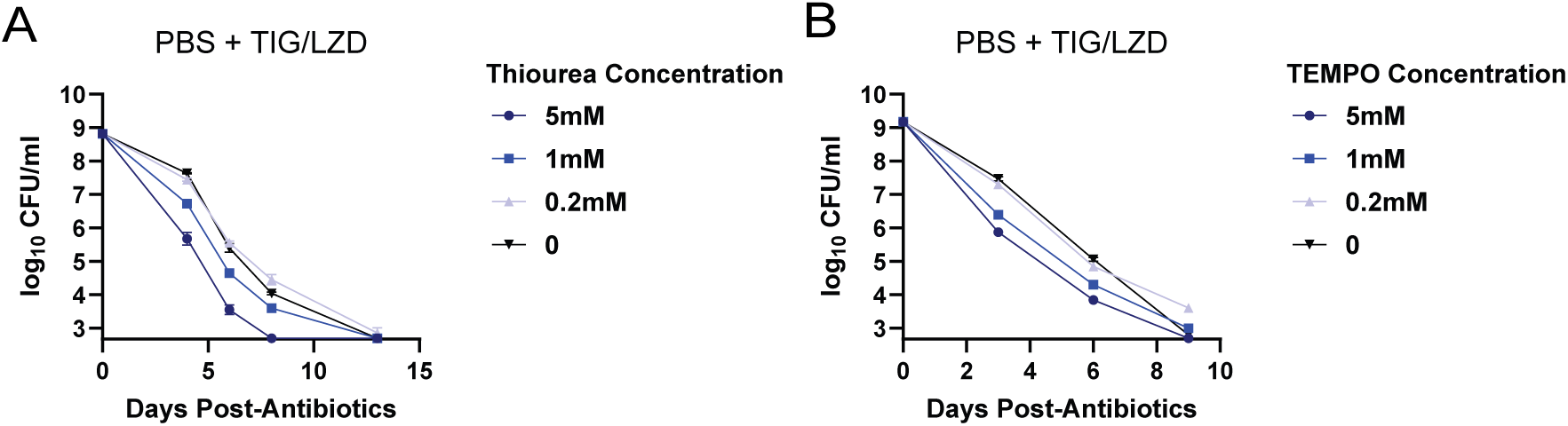
Effect of ROS scavengers. *ΔkatG* cells were starved in PBS for 48h then treated with TIG/LZD and the indicated concentration of (A) thiourea or (B) TEMPO and surviving CFU over time were measured. Representative data from 2 independent experiments are shown.

Supplemental Table 1. Tn-Seq results

Supplemental Table 2. DAVID functional pathway analysis

Supplemental Table 3. Oligonucleotide sequences

Supplemental Table 4. 16s sequencing data of clinical strains

Supplemental Table 5. MIC Values

## Notes

### Summary of Updates

Text changes for clarity and additional references.

## REFERENCES

1. Stevens, D.L., A.L. Bisno, H.F. Chambers, E.P. Dellinger, E.J. Goldstein, S.L. Gorbach, J.V. Hirschmann, S.L. Kaplan, J.G. Montoya, and J.C. Wade, Practice guidelines for the diagnosis and management of skin and soft tissue infections: 2014 update by the Infectious Diseases Society of America. Clin Infect Dis, 2014. 59(2): p. e10–52.

2. Metersky, M.L. and A.C. Kalil, Management of ventilator-associated pneumonia: guidelines. Clinics in chest medicine, 2018. 39(4): p. 797–808.

3. Nahid, P., S.E. Dorman, N. Alipanah, P.M. Barry, J.L. Brozek, A. Cattamanchi, L.H. Chaisson, R.E. Chaisson, C.L. Daley, and M. Grzemska, Official American thoracic society/centers for disease control and prevention/infectious diseases society of America clinical practice guidelines: treatment of drug-susceptible tuberculosis. Clinical infectious diseases, 2016. 63(7): p. e147–e195.

4. Griffith, D.E. and C.L. Daley, Treatment of Mycobacterium abscessus Pulmonary Disease. CHEST, 2022. 161(1): p. 64–75.

5. Griffith, D.E., T. Aksamit, B.A. Brown-Elliott, A. Catanzaro, C. Daley, F. Gordin, S.M. Holland, R. Horsburgh, G. Huitt, and M.F. Iademarco, An official ATS/IDSA statement: diagnosis, treatment, and prevention of nontuberculous mycobacterial diseases. American journal of respiratory and critical care medicine, 2007. 175(4): p. 367–416.

6. Meylan, S., I.W. Andrews, and J.J. Collins, Targeting Antibiotic Tolerance, Pathogen by Pathogen. Cell, 2018. 172(6): p. 1228–1238.

7. Namugenyi, S.B., A.M. Aagesen, S.R. Elliott, and A.D. Tischler, Mycobacterium tuberculosis PhoY Proteins Promote Persister Formation by Mediating Pst/SenX3-RegX3 Phosphate Sensing. mBio, 2017. 8(4): p. e00494–17.

8. Liu, Y., S. Tan, L. Huang, R.B. Abramovitch, K.H. Rohde, M.D. Zimmerman, C. Chen, V. Dartois, B.C. VanderVen, and D.G. Russell, Immune activation of the host cell induces drug tolerance in Mycobacterium tuberculosis both in vitro and in vivo. Journal of Experimental Medicine, 2016. 213(5): p. 809–825.

9. Gold, B. and C. Nathan, Targeting Phenotypically Tolerant Mycobacterium tuberculosis. Microbiol Spectr, 2017. 5(1).

10. Bigger, J., Treatment of Staphyloeoeeal Infections with Penicillin by Intermittent Sterilisation. Lancet, 1944: p. 497–500.

11. Ronneau, S., P.W.S. Hill, and S. Helaine, Antibiotic persistence and tolerance: not just one and the same. Current opinion in microbiology, 2021. 64: p. 76–81.

12. Dhar, N. and J.D. McKinney, Microbial phenotypic heterogeneity and antibiotic tolerance. Current opinion in microbiology, 2007. 10(1): p. 30–38.

13. Grant, S.S., B.B. Kaufmann, N.S. Chand, N. Haseley, and D.T. Hung, Eradication of bacterial persisters with antibiotic-generated hydroxyl radicals. Proceedings of the National Academy of Sciences, 2012. 109(30): p. 12147–12152.

14. Baker, J.J. and R.B. Abramovitch, Genetic and metabolic regulation of Mycobacterium tuberculosis acid growth arrest. Scientific Reports, 2018. 8(1).

15. Saito, K., T. Warrier, S. Somersan-Karakaya, L. Kaminski, J. Mi, X. Jiang, S. Park, K. Shigyo, B. Gold, and J. Roberts, Rifamycin action on RNA polymerase in antibiotic-tolerant Mycobacterium tuberculosis results in differentially detectable populations. Proceedings of the National Academy of Sciences, 2017. 114(24): p. E4832–E4840.

16. Hipolito, V.E.B., E. Ospina-Escobar, and R.J. Botelho, Lysosome remodelling and adaptation during phagocyte activation. Cell Microbiol, 2018. 20(4).

17. Sukumar, N., S. Tan, B.B. Aldridge, and D.G. Russell, Exploitation of Mycobacterium tuberculosis reporter strains to probe the impact of vaccination at sites of infection. PLoS Pathog, 2014. 10(9): p. e1004394.

18. Schumacher, M.A., K.M. Piro, W. Xu, S. Hansen, K. Lewis, and R.G. Brennan, Molecular Mechanisms of HipA-Mediated Multidrug Tolerance and Its Neutralization by HipB. Science, 2009. 323(5912): p. 396–401.

19. Kusser, W. and E.E. Ishiguro, Suppression of mutations conferring penicillin tolerance by interference with the stringent control mechanism of Escherichia coli. Journal of Bacteriology, 1987. 169(9): p. 4396–4398.

20. Rodionov, D.G. and E.E. Ishiguro, Direct correlation between overproduction of guanosine 3’,5’-bispyrophosphate (ppGpp) and penicillin tolerance in Escherichia coli. Journal of Bacteriology, 1995. 177(15): p. 4224–4229.

21. Bokinsky, G., E.E.K. Baidoo, S. Akella, H. Burd, D. Weaver, J. Alonso-Gutierrez, H. García-Martín, T.S. Lee, and J.D. Keasling, HipA-Triggered Growth Arrest and β-Lactam Tolerance in Escherichia coli Are Mediated by RelA-Dependent ppGpp Synthesis. Journal of Bacteriology, 2013. 195(14): p. 3173–3182.

22. Korch, S.B., T.A. Henderson, and T.M. Hill, Characterization of the hipA7 allele of Escherichia coli and evidence that high persistence is governed by (p)ppGpp synthesis. Mol Microbiol, 2003. 50(4): p. 1199–213.

23. Harms, A., C. Fino, A. Sørensen Michael, S. Semsey, and K. Gerdes, Prophages and Growth Dynamics Confound Experimental Results with Antibiotic-Tolerant Persister Cells. mBio, 2017. 8(6): p. 10.1128/mbio.01964-17.

24. Wong, F., S. Wilson, R. Helbig, S. Hegde, O. Aftenieva, H. Zheng, C. Liu, T. Pilizota, E.C. Garner, A. Amir, and L.D. Renner, Understanding Beta-Lactam-Induced Lysis at the Single-Cell Level. Frontiers in Microbiology, 2021. 12.

25. Kohanski, M.A., D.J. Dwyer, B. Hayete, C.A. Lawrence, and J.J. Collins, A common mechanism of cellular death induced by bactericidal antibiotics. Cell, 2007. 130(5): p. 797–810.

26. Dwyer, D.J., P.A. Belenky, J.H. Yang, I.C. MacDonald, J.D. Martell, N. Takahashi, C.T.Y. Chan, M.A. Lobritz, D. Braff, E.G. Schwarz, J.D. Ye, M. Pati, M. Vercruysse, P.S. Ralifo,

27. K.R. Allison, A.S. Khalil, A.Y. Ting, G.C. Walker, and J.J. Collins, Antibiotics induce redox-related physiological alterations as part of their lethality. Proceedings of the National Academy of Sciences, 2014. 111(20): p. E2100–E2109.

27. Zeng, J., Y. Hong, N. Zhao, Q. Liu, W. Zhu, L. Xiao, W. Wang, M. Chen, S. Hong, L. Wu, Y. Xue, D. Wang, J. Niu, K. Drlica, and X. Zhao, A broadly applicable, stress-mediated bacterial death pathway regulated by the phosphotransferase system (PTS) and the cAMP-Crp cascade. Proc Natl Acad Sci U S A, 2022. 119(23): p. e2118566119.

28. Shee, S., S. Singh, A. Tripathi, C. Thakur, T.A. Kumar, M. Das, V. Yadav, S. Kohli, R.S. Rajmani, N. Chandra, H. Chakrapani, K. Drlica, and A. Singh, Moxifloxacin-Mediated Killing of Mycobacterium tuberculosis Involves Respiratory Downshift, Reductive Stress, and Accumulation of Reactive Oxygen Species. Antimicrob Agents Chemother, 2022. 66(9): p. e0059222.

29. Saito, K., S. Mishra, T. Warrier, N. Cicchetti, J. Mi, E. Weber, X. Jiang, J. Roberts, A. Gouzy, E. Kaplan, C.D. Brown, B. Gold, and C. Nathan, Oxidative damage and delayed replication allow viable Mycobacterium tuberculosis to go undetected. Sci Transl Med, 2021. 13(621): p. eabg2612.

30. Vilchèze, C., T. Hartman, B. Weinrick, P. Jain, T.R. Weisbrod, L.W. Leung, J.S. Freundlich, and W.R. Jacobs, Jr., Enhanced respiration prevents drug tolerance and drug resistance in Mycobacterium tuberculosis. Proc Natl Acad Sci U S A, 2017. 114(17): p. 4495–4500.

31. Augusto Cesar, H.-S., J.P. Brian, B. Joseph, and C.B. Cara, Mycobacterium abscessus Cells Have Altered Antibiotic Tolerance and Surface Glycolipids in Artificial Cystic Fibrosis Sputum Medium. Antimicrobial Agents and Chemotherapy, 2019. 63(7): p. e02488–18.

32. Li, S., N.C. Poulton, J.S. Chang, Z.A. Azadian, M.A. DeJesus, N. Ruecker, M.D. Zimmerman, K.A. Eckartt, B. Bosch, and C.A. Engelhart, CRISPRi chemical genetics and comparative genomics identify genes mediating drug potency in Mycobacterium tuberculosis. Nature microbiology, 2022. 7(6): p. 766–779.

33. Carey, A.F., J.M. Rock, I.V. Krieger, M.R. Chase, M. Fernandez-Suarez, S. Gagneux, J.C. Sacchettini, T.R. Ioerger, and S.M. Fortune, TnSeq of Mycobacterium tuberculosis clinical isolates reveals strain-specific antibiotic liabilities. PLoS pathogens, 2018. 14(3): p. e1006939.

34. Rodriguez, R., N. Campbell-Kruger, J. Gonzalez Camba, J. Berude, R. Fetterman, and S. Stanley, MarR-Dependent Transcriptional Regulation of mmpSL5 Induces Ethionamide Resistance in Mycobacterium abscessus. Antimicrobial Agents and Chemotherapy, 2023. 67(4): p. e01350–22.

35. Kreutzfeldt, K.M., R.S. Jansen, T.E. Hartman, A. Gouzy, R. Wang, I.V. Krieger, M.D. Zimmerman, M. Gengenbacher, J.P. Sarathy, M. Xie, V. Dartois, J.C. Sacchettini, K.Y. Rhee, D. Schnappinger, and S. Ehrt, CinA mediates multidrug tolerance in Mycobacterium tuberculosis. Nat Commun, 2022. 13(1): p. 2203.

36. Bellerose, M.M., S.-H. Baek, C.-C. Huang, C.E. Moss, E.-I. Koh, M.K. Proulx, C.M. Smith, R.E. Baker, J.S. Lee, S. Eum, S.J. Shin, S.-N. Cho, M. Murray, and C.M. Sassetti, Common Variants in the Glycerol Kinase Gene Reduce Tuberculosis Drug Efficacy. mBio, 2019. 10(4): p. e00663–19.

37. Kumar, K., C.L. Daley, D.E. Griffith, and M.R. Loebinger, Management of Mycobacterium avium complex and Mycobacterium abscessus pulmonary disease: therapeutic advances and emerging treatments. European respiratory review, 2022. 31(163).

38. Yam, Y.-K., N. Alvarez, M.-L. Go, and T. Dick, Extreme Drug Tolerance of Mycobacterium abscessus “Persisters”. Frontiers in Microbiology, 2020. 11.

39. Berube, B.J., L. Castro, D. Russell, Y. Ovechkina, and T. Parish, Novel Screen to Assess Bactericidal Activity of Compounds Against Non-replicating Mycobacterium abscessus. Front Microbiol, 2018. 9: p. 2417.

40. Lee, J., N. Ammerman, A. Agarwal, M. Naji, S.Y. Li, and E. Nuermberger, Differential In Vitro Activities of Individual Drugs and Bedaquiline-Rifabutin Combinations against Actively Multiplying and Nutrient-Starved Mycobacterium abscessus. Antimicrob Agents Chemother, 2021. 65(2).

41. Betts, J.C., P.T. Lukey, L.C. Robb, R.A. McAdam, and K. Duncan, Evaluation of a nutrient starvation model of Mycobacterium tuberculosis persistence by gene and protein expression profiling. Molecular microbiology, 2002. 43(3): p. 717–731.

42. DeJesus, M.A., C. Ambadipudi, R. Baker, C. Sassetti, and T.R. Ioerger, TRANSIT-a software tool for Himar1 TnSeq analysis. PLoS computational biology, 2015. 11(10): p. e1004401.

43. Huang da, W., B.T. Sherman, and R.A. Lempicki, Systematic and integrative analysis of large gene lists using DAVID bioinformatics resources. Nat Protoc, 2009. 4(1): p. 44–57.

44. Darwin, K.H., S. Ehrt, J.-C. Gutierrez-Ramos, N. Weich, and C.F. Nathan, The proteasome of Mycobacterium tuberculosis is required for resistance to nitric oxide. Science, 2003. 302(5652): p. 1963-1966.

45. Shi, H., R. Zhang, L. Lan, Z. Chen, and J. Kan, Zinc mediates resuscitation of lactic acid-injured Escherichia coli by relieving oxidative stress. Journal of Applied Microbiology, 2019. 127(6): p. 1741–1750.

46. Hohle, T.H. and M.R. O’Brian, The mntH gene encodes the major Mn(2+) transporter in Bradyrhizobium japonicum and is regulated by manganese via the Fur protein. Mol Microbiol, 2009. 72(2): p. 399–409.

47. Murphy, K.C., S.J. Nelson, S. Nambi, K. Papavinasasundaram, C.E. Baer, and C.M. Sassetti, ORBIT: a new paradigm for genetic engineering of mycobacterial chromosomes. mBio, 2018. 9(6): p. e01467–18.

48. McBee, M.E., Y.H. Chionh, M.L. Sharaf, P. Ho, M.W.L. Cai, and P.C. Dedon, Production of superoxide in bacteria is stress- and cell state-dependent: A gating-optimized flow cytometry method that minimizes ROS measurement artifacts with fluorescent dyes. Frontiers in Microbiology, 2017. 8(MAR).

49. Wayne, L.G. and L.G. Hayes, An In Vitro Model for Sequential Study of Shiftdown of Mycobacterium tuberculosis through Two Stages of Nonreplicating Persistence. Infection and Immunity, 1996. 64(6): p. 2062–2069.

50. Winterbourn, C.C., Toxicity of iron and hydrogen peroxide: the Fenton reaction. Toxicol Lett, 1995. 82-83: p. 969-74.

51. Moyed, H.S. and K.P. Bertrand, hipA, a newly recognized gene of Escherichia coli K-12 that affects frequency of persistence after inhibition of murein synthesis. Journal of Bacteriology, 1983. 155(2): p. 768–775.

52. Viducic, D., T. Ono, K. Murakami, H. Susilowati, S. Kayama, K. Hirota, and Y. Miyake, Functional analysis of spoT, relA and dksA genes on quinolone tolerance in Pseudomonas aeruginosa under nongrowing condition. Microbiol Immunol, 2006. 50(4): p. 349–57.

53. Geiger, T., B. Kästle, F.L. Gratani, C. Goerke, and C. Wolz, Two small (p)ppGpp synthases in Staphylococcus aureus mediate tolerance against cell envelope stress conditions. J Bacteriol, 2014. 196(4): p. 894–902.

54. Dutta, N.K., L.G. Klinkenberg, M.J. Vazquez, D. Segura-Carro, G. Colmenarejo, F. Ramon, B. Rodriguez-Miquel, L. Mata-Cantero, E. Porras-De Francisco, Y.M. Chuang, H. Rubin, J.J. Lee, H. Eoh, J.S. Bader, E. Perez-Herran, A. Mendoza-Losana, and P.C. Karakousis, Inhibiting the stringent response blocks Mycobacterium tuberculosis entry into quiescence and reduces persistence. Sci Adv, 2019. 5(3): p. eaav2104.

55. Bhaskar, A., C. De Piano, E. Gelman, J.D. McKinney, and N. Dhar, Elucidating the role of (p) ppGpp in mycobacterial persistence against antibiotics. IUBMB life, 2018. 70(9): p. 836–844.

56. Wu, N., L. He, P. Cui, W. Wang, Y. Yuan, S. Liu, T. Xu, S. Zhang, J. Wu, W. Zhang, and Y. Zhang, Ranking of persister genes in the same Escherichia coli genetic background demonstrates varying importance of individual persister genes in tolerance to different antibiotics. Front Microbiol, 2015. 6: p. 1003.

57. Zeng, J., Y. Hong, N. Zhao, Q. Liu, W. Zhu, L. Xiao, W. Wang, M. Chen, S. Hong, L. Wu, Y. Xue, D. Wang, J. Niu, K. Drlica, and X. Zhao, A broadly applicable, stress-mediated bacterial death pathway regulated by the phosphotransferase system (PTS) and the cAMP-Crp cascade. Proceedings of the National Academy of Sciences, 2022. 119(23): p. e2118566119.

58. Hunt-Serracín, A.C., M.I. Kazi, J.M. Boll, and C.C. Boutte, In Mycobacterium abscessus, the Stringent Factor Rel Regulates Metabolism but Is Not the Only (p)ppGpp Synthase. J Bacteriol, 2022. 204(2): p. e0043421.

59. Liu, Y. and J.A. Imlay, Cell death from antibiotics without the involvement of reactive oxygen species. Science, 2013. 339(6124): p. 1210-3.

60. Keren, I., Y. Wu, J. Inocencio, L.R. Mulcahy, and K. Lewis, Killing by bactericidal antibiotics does not depend on reactive oxygen species. Science, 2013. 339(6124): p. 1213-6.

61. Mahoney, T.F. and T.J. Silhavy, The Cpx stress response confers resistance to some, but not all, bactericidal antibiotics. J Bacteriol, 2013. 195(9): p. 1869–74.

62. Ezraty, B., A. Vergnes, M. Banzhaf, Y. Duverger, A. Huguenot, A.R. Brochado, S.Y. Su, L. Espinosa, L. Loiseau, B. Py, A. Typas, and F. Barras, Fe-S cluster biosynthesis controls uptake of aminoglycosides in a ROS-less death pathway. Science, 2013. 340(6140): p. 1583-7.

63. Goswami, M., S.H. Mangoli, and N. Jawali, Involvement of Reactive Oxygen Species in the Action of Ciprofloxacin against Escherichia coli. Antimicrobial Agents and Chemotherapy, 2006. 50(3): p. 949–954.

64. Wang, X. and X. Zhao, Contribution of oxidative damage to antimicrobial lethality. Antimicrob Agents Chemother, 2009. 53(4): p. 1395–402.

65. Hong, Y., Q. Li, Q. Gao, J. Xie, H. Huang, K. Drlica, and X. Zhao, Reactive oxygen species play a dominant role in all pathways of rapid quinolone-mediated killing. J Antimicrob Chemother, 2020. 75(3): p. 576–585.

66. Snapper, S.B., L. Lugosi, A. Jekkel, R.E. Melton, T. Kieser, B.R. Bloom, and W.R. Jacobs, Jr., Lysogeny and transformation in mycobacteria: stable expression of foreign genes. Proc Natl Acad Sci U S A, 1988. 85(18): p. 6987–91.

